# gbdraw: a genome diagram generator for microbes and organelles

**DOI:** 10.64898/2026.04.07.716863

**Authors:** Satoshi Kawato

**Author notes:** Corresponding author: Satoshi Kawato.

## Abstract

**Motivation:** Generating graphical diagrams of microbial and organellar genomes is a common and essential task in bioinformatics. Existing tools often present a trade-off; while powerful programming libraries that require coding skills, graphical applications require server processing or local installation with complex dependency. This highlights the need for a tool that offers both programmatic control for batch processing and graphical accessibility for ease of use.

**Results:** To fill this gap, I developed gbdraw, a web application that generates circular and linear genome diagrams from self-contained GenBank or DDBJ files or combinations of GFF3 annotation and FASTA sequence files. Its core functions include visualizing annotated features, plotting GC content/skew tracks, and optionally generating pairwise sequence comparisons for comparative genomics. It is available as both a GUI web application and a command-line utility. Unlike existing web-based tools that require data upload to a remote server, gbdraw operates entirely within the user’s web browser. This serverless architecture ensures that sensitive sequence data never leaves the local machine, providing a secure environment for visualizing unpublished genomic data.

**Availability and Implementation:** gbdraw is implemented in Python 3 (version 3.10+) and is freely available under the MIT license. The web app is available at https://gbdraw.app/. Source code and documentation are available at https://github.com/satoshikawato/gbdraw. The local version can be installed from the Bioconda channel using a conda-compatible package manager.

## Introduction

Visualizing genome sequences is essential for understanding gene organization and evolutionary relationships. A diverse ecosystem of software tools has emerged to meet this demand, each catering to different needs. For interactive, hands-on exploration, desktop applications like EasyFig (Sullivan, Petty, and Beatson 2011), Artemis Comparison Tool (Carver *et al*. 2008), SnapGene Viewer (https://www.snapgene.com/), and GenomeMatcher (Ohtsubo *et al*. 2008) offer rich graphical interfaces. Command-line tools like Genovi (Cumsille *et al*. 2023) and lovis4u (Egorov and Atkinson 2025) offer specialized annotation and visualization suites for prokaryotes and bacteriophages, respectively. Comprehensive web-based platforms such as Proksee (Grant *et al*. 2023) and Bakta (Schwengers *et al*. 2021) provide one-stop pipelines that guide users from annotation to visualization, while specialized tools like OGDRAW (Greiner, Lehwark, and Bock 2019) are tailored to the specific conventions of organellar genomics. Clinker (Gilchrist and Chooi 2021) excels in gene cluster comparison figures. Programming libraries like Circos (Krzywinski *et al*. 2009) in Perl, pyGenomeViz (https://github.com/moshi4/pyGenomeViz) in Python, as well as gggenomes (Hackl *et al*. 2024) and geneviewer (https://nvelden.github.io/geneviewer/) in R enable deep customization and integration into bioinformatic workflows.

Despite this rich ecosystem offering powerful options, a practical gap remains for many researchers. Programming libraries offer extensive control and can produce high-quality graphics, but achieving an optimal output can require a deeper understanding of the library’s parameters and intensive coding. While many GUI-based tools do offer customization options, server-hosted services require online connection and remote processing of user’s sequence data. Local tools often require additional dependencies, e.g. BLAST, which could be cumbersome for beginners. Moreover, most tools cover either circular or linear topology only, which necessitates switching between multiple tools to visualize different aspects of their datasets.

Here I present gbdraw, a genome diagram generator designed to bridge this gap by balancing accessibility and customizability. gbdraw offers both an intuitive web application for rapid, code-free visualization and a command-line interface (CLI) suitable for scripting. A defining feature of the web application is its serverless architecture. By executing all processes locally in the browser, gbdraw ensures that sensitive sequence data never leaves the user’s machine, providing a secure environment without the need for local software installation.

## Implementation and Features

### Design and implementation

gbdraw is a Python-based tool that accepts annotated genomes in GenBank/DDBJ format or pairs of GFF3 and FASTA files as input. gbdraw is accessible via two interfaces; the web application (https://gbdraw.app/) provides an interactive interface where users can upload files, adjust parameters via widgets, and download the resulting images, requiring no local installation or programming experience. The CLI is designed for integration into scripts and automated pipelines.

gbdraw uses Biopython (Cock *et al*. 2009) for parsing input files, pandas (The pandas development team 2020) for data handling, and svgwrite (https://github.com/mozman/svgwrite) for generating SVG graphics. The web application utilizes Pyodide (Chatham *et al*. 2026) to run the core Python logic directly within the web browser. For image conversion, the CLI version uses CairoSVG (https://cairosvg.org/) for PNG/PDF generation; the web application uses the HTML5 Canvas API to generate PNG images, while relying on JavaScript libraries jsPDF (https://github.com/MrRio/jsPDF) and svg2pdf.js (https://github.com/yWorks/svg2pdf.js/) for SVG to PDF conversion.

The web application leverages Pyodide (Chatham *et al*. 2026) to execute the core Python logic directly within the user’s web browser via WebAssembly. This approach eliminates the latency and privacy concerns associated with data transfer to a backend server, allowing the application to utilize local computing resources for immediate rendering.

A distinctive feature of the gbdraw web application is the integration of LOSAT (https://github.com/satoshikawato/LOSAT), a homology search engine designed for the browser. LOSAT enables on-the-fly generation of pairwise genomic comparisons (equivalent to TBLASTX or BLASTN) without requiring the user to install the BLAST+ suite locally. For large-scale comparisons where browser-based search might be performance-limited, the tool remains flexible by allowing the import of pre-calculated BLAST output files.

gbdraw allows users to export the entire state of their visualization session as a .json file. This file captures all parameters, track configurations, and custom edits, enabling researchers to resume their work or share the exact visualization settings with collaborators.

gbdraw can render genome diagrams in both circular and linear topologies (Figures 1 and 2), with sizes ranging from eukaryotic organelles (Figure 1A, B) to prokaryotic chromosomes (Figure 1C). In circular mode, genes are rendered on concentric tracks, and GC content/skew tracks can be added to help identify genomic features like the origin of replication and recent episodes of genome reshuffling. Multiple records can be displayed on a single canvas to give a comprehensive view of multi-replicon genomes, e.g. *Vibrio* spp. (Figure 1C).

**Figure 1.**
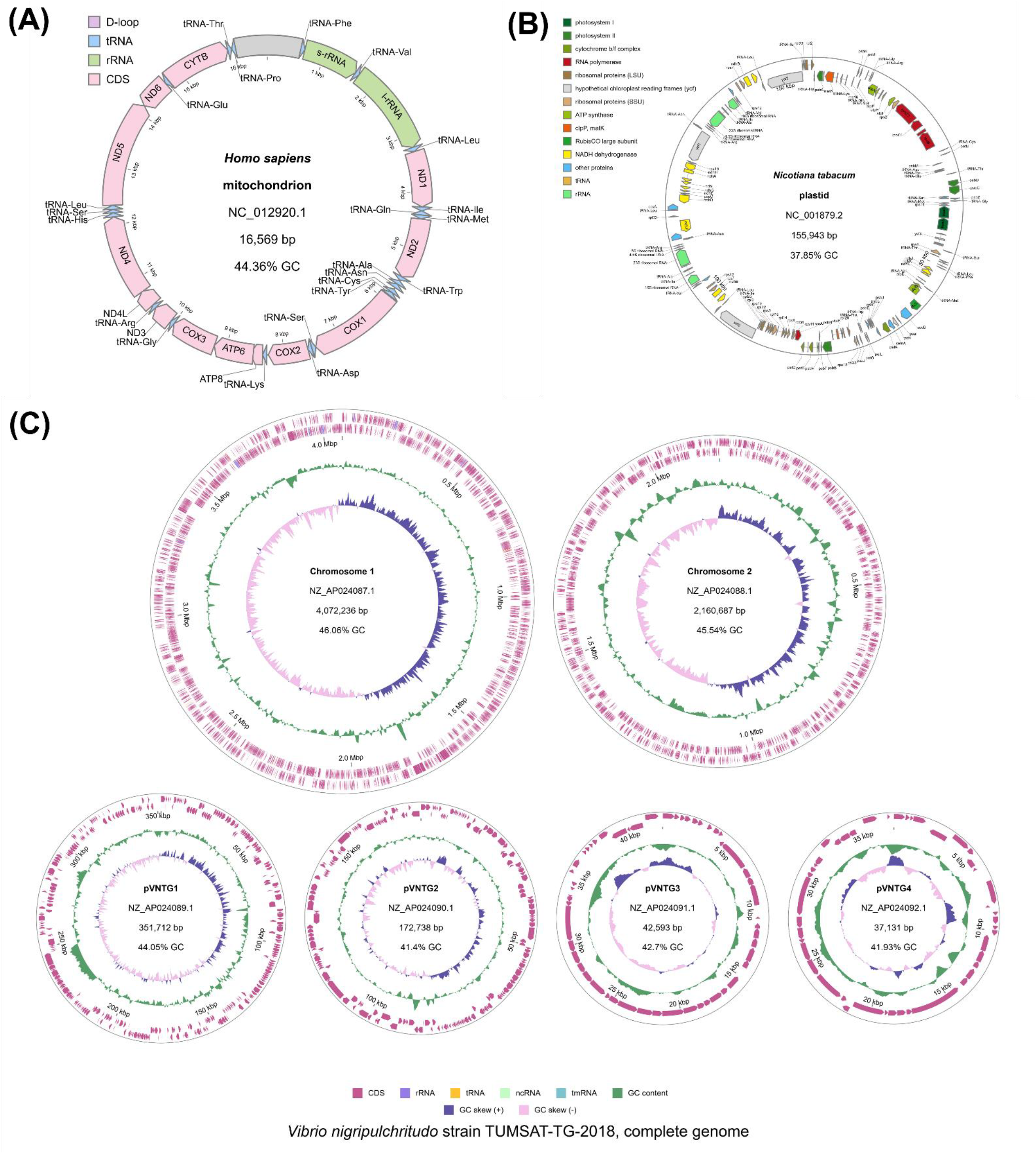
gbdraw circular diagrams. (A) Circular diagram of the human mitochondrial DNA (NC_012920.1). “soft_pastel” palette was used. (B) Circular diagram of the tobacco plastid genome (NC_001879.1). Color scheme deferred to that of OGDRAW. (C) Circular diagram of *Vibrio nigripulchritudo* TUMSAT-TG-2018. A total of six replicons are arranged in order of size. Diagram radii do not reflect the actual size difference.

**Figure 2.**
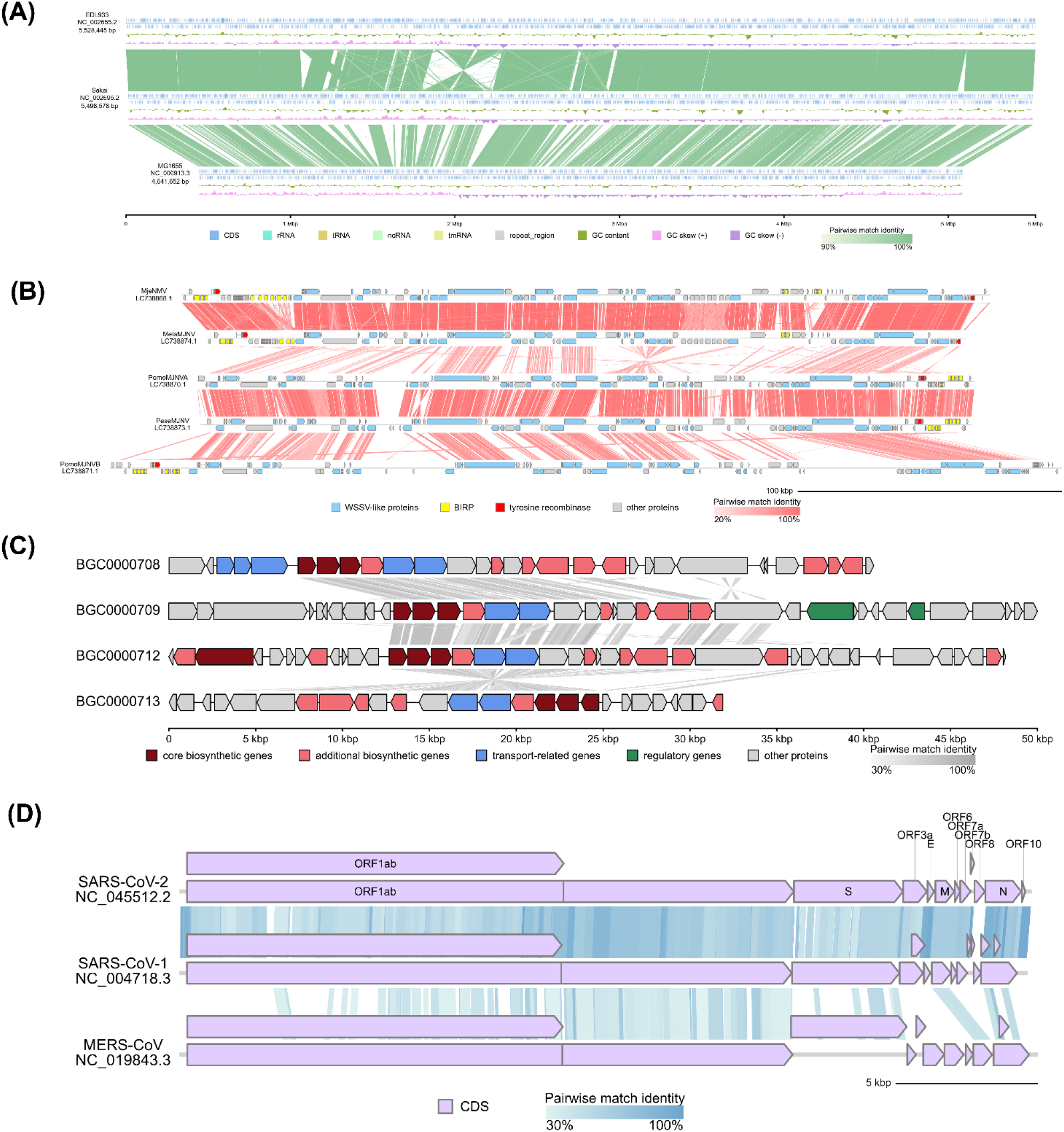
gbdraw linear diagrams. (A) Linear diagram of *Escherichia coli* chromosomes. “ajisai” palette was used. Genome diagrams are linked by LOSATN pairwise matches. (B) Linear diagram of majaniviruses, a group of endogenous DNA virus colonizing penaeid shrimps. Genome diagrams are linked by TBLASTX pairwise matches. (C) Linear diagram of lividomycin-like aminoglycoside biosynthetic gene clusters. Records are connected by ribbons representing LOSATX pairwise matches. Color schemes for gene categories are adapted from the MIBiG repository (https://mibig.secondarymetabolites.org/repository). (D) Linear diagram of betacoronaviruses SARS-CoV-2, SARS-CoV-1, and MERS-CoV. The records are connected by ribbons TLOSATX pairwise matches.

The linear mode is suitable for visualizing synteny and structural rearrangements between multiple genomes or genomic segments (Figure 2). Although gbdraw is not fit for visualizing eukaryotic whole genomes, it can be used to visualize regions of interest. gbdraw natively handles features with introns, which is crucial for certain eukaryotic viruses (Figure 2B). For comparative genomics, pairwise alignment data can be generated on-the-fly using the built-in LOSAT engine without requiring a local BLAST+ installation. For computationally expensive tasks, such as TBLASTX searches of large bacterial genomes, gbdraw remains flexible by allowing the import of pre-calculated BLAST output files in tab- or comma-separated formats.

The visual appearance of diagrams is highly customizable. The tool includes 55 built-in color palettes that can be applied globally. For more specific control, users can define colors for specific feature types (e.g., CDS, tRNA) or for individual features. Labels can be granularly controlled. Users can define the priority of feature qualifiers to be used for labels (e.g., prefer ‘gene’ over ‘product’), adjust font sizes, and provide blacklist or whitelist files to focus the audience’s attention to specific features. Features and labels can be edited directly on the webapp, while on CLI, the user can provide tab-separated value (TSV) files via command-line options.

## Discussion

gbdraw is a practical tool for microbial and organellar genome visualization that balances accessibility and automation. Its dual-interface design serves both computational and experimental biologists. The development of gbdraw was motivated by the visualization needs of our previous research on bacteria and large double-stranded DNA viruses. The core methods were initially created as a series of custom scripts to generate figures for those works, and have since been refined, generalized, and packaged into the user-friendly tool presented here.

gbdraw provides a practical solution for one of the most common and essential visualization steps in modern genomics research. Future development will focus on expanding the repertoire of visualization options, such as incorporating additional information tracks and further streamlining the user customization process.

## Acknowledgements

I thank Yamato Matsuoka for his valuable advice on publishing gbdraw to the Bioconda channel.

## Data availability

gbdraw is open-source software available under the MIT license. The source code and documentation are available on GitHub at https://github.com/satoshikawato/gbdraw. The tool can be installed via the Bioconda channel. The web app is available at https://gbdraw.app/. The source code of LOSAT is available at https://github.com/satoshikawato/LOSAT.

